# Simultaneous dual-color calcium imaging in freely-behaving mice

**DOI:** 10.1101/2024.07.03.601770

**Authors:** Zhe Dong, Yu Feng, Keziah Diego, Austin M. Baggetta, Brian M. Sweis, Zachary T. Pennington, Sophia I. Lamsifer, Yosif Zaki, Federico Sangiuliano, Paul A. Philipsberg, Denisse Morales-Rodriguez, Daniel Kircher, Paul Slesinger, Tristan Shuman, Daniel Aharoni, Denise J. Cai

## Abstract

Miniaturized fluorescence microscopes (miniscopes) enable imaging of calcium events from a large population of neurons in freely behaving animals. Traditionally, miniscopes have only been able to record from a single fluorescence wavelength. Here, we present a new open-source dual-channel Miniscope that simultaneously records two wavelengths in freely behaving animals. To enable simultaneous acquisition of two fluorescent wavelengths, we incorporated two CMOS sensors into a single Miniscope. To validate our dual-channel Miniscope, we imaged hippocampal CA1 region that co-expressed a dynamic calcium indicator (GCaMP) and a static nuclear signal (tdTomato) while mice ran on a linear track. Our results suggest that, even when neurons were registered across days using tdTomato signals, hippocampal spatial coding changes over time. In conclusion, our novel dual-channel Miniscope enables imaging of two fluorescence wavelengths with minimal crosstalk between the two channels, opening the doors to a multitude of new experimental possibilities.

**Teaser:** Novel open-source dual-channel Miniscope that simultaneously records two wavelengths with minimal crosstalk in freely behaving animals.

## Introduction

Recent years have seen rapid growth in the development and dissemination of open-source miniature microscopes (miniscopes) [1,2,3]. The miniscope allows circuit investigation of large populations of neurons across days in small animals such as rodents [4,5,6,7,8,9,10,11], birds [12], and bats [13]. Since miniscopes are lightweight enough to be carried by animals while freely behaving, a broad range of rich behaviors can be studied including song vocalization [12], social interaction [6,10], spatial navigation [4], memory integration and updating [5]. The open-source UCLA Miniscope project is an impactful open-source initiative in the neuroscience community, with over 700 labs building and using Miniscopes to carry out independent research [1,2]. Since the initial release, the UCLA Miniscope has iterated through several versions. The latest version of the UCLA Miniscope (v4) features a highly sensitive PYTHON480 CMOS sensor, a large 1mm x 1mm field of view, fully achromatic optics, and electrowetting focus adjustment of ±200µm (https://github.com/Aharoni-Lab/Miniscope-v4) [5]. The open-source nature of the UCLA Miniscope also facilitates customization and innovations on top of the original design. Several alternative designs have improved imaging sensitivity, provided a more compact microscope size, larger field of view, and lighter weight, which collectively enables imaging across multiple brain regions and better overall quality [14,15,16,17,18,19,20]. In addition, wireless versions of the Miniscope allow animals to travel in a large environment and enable calcium imaging in behaving songbirds and bats [11,21]. Furthermore, 3D imaging using Miniscopes is possible with both computational imaging and traditional optical imaging techniques [22,23,24,25]. In summary, with these recent technological advancements, it is prime time to innovate on top of the latest hardware from the UCLA Miniscope project and further expand the functionality of Miniscopes to answer novel research questions.

One of the most common use cases for Miniscopes is to image genetically encoded calcium indicators like GCaMP, which allows measurement of neural population activity at single-cell resolution [2]. However, with the advancement of new genetic tools, it is beneficial to combine two or more fluorescent reporters to answer questions about the nervous system in unprecedented ways. There is a growing need to image two or more fluorescent reporters simultaneously at single-cell resolution. Multiple scenarios could benefit from dual-channel imaging. For example, one can image a GCaMP signal together with a static neuronal marker in the other channel. The static neuronal signal can improve motion correction within a recording session, image registration across recording sessions, and identification of neurons that are inactive during the recording. In addition, by using cell-type or activity-dependent markers, researchers may use the additional channel to differentiate cell types and characterize differences in neural activity specific to the genetic and activation profiles of neural ensembles (e.g., a static neuronal marker might identify GABAergic neurons while GCaMP is used to record from all neurons). Similarly, simultaneous imaging of two dynamic fluorescence signals across the two channels could shine light on how different neural populations coordinate and function together. For example, one may use GCaMP and RCaMP (a red-shifted calcium indicator) to mark excitatory and inhibitory neurons, respectively, and subsequently study the population dynamics of these two distinct populations. In summary, developing a Miniscope that is capable of dual-channel imaging in behaving animals will fill a significant gap in current neural imaging technologies.

One important application for a dual-channel Miniscope is to use the additional channel to image a static neuronal marker labeling all neurons in the field of view, which allows researchers to track a neural population over time. Despite recent developments in registration algorithms [26], tracking the same population of cells across long periods of time (e.g., weeks to months) remains a challenge in calcium imaging, due to the dynamic nature of GCaMP signals (i.e., not all cells are active on a given recording session). The ability to track a neural population over time is particularly important when studying the stability of neural ensembles (e.g., hippocampal spatial representations). Recently, it has been reported that even when animals performed the same spatial navigation task in the same environment, the hippocampal spatial representation changed over time (i.e., representational drift) [4,27]. This change is often described as a decrease of spatial representation stability (such as reactivation rate of place cells and population vector correlation) over time. However, since the imaging field of view may slightly change across days in behaving animals, it was previously unclear how much the instability of the field of view contributed to the instabilities in reactivation rate and population vector correlation. For instance, if a cell that appears in one session lies near the edge of the field of view (in the x-y plane or z-direction), imaging this same cell in a subsequent session with a slightly changed field of view may result in a significant change in signal level, since the cell may no longer be in focus or may be completely outside the field of view. In other words, instability in the field of view alone could result in a “false” change in reactivation rate or population vector correlation that should not be attributed to biological changes in neural activity patterns. As a result, researchers often have to fall back to more conservative heuristics when measuring the change in hippocampal spatial representation. One common approach is to include only the neurons that can be registered using GCaMP signal throughout the experiment, which limits the analysis to the population of neurons that were always active over time, yielding a partial view of how the hippocampal spatial representation changes. In contrast, collecting a static landmark signal for each cell together with the GCaMP signal would allow us to detect and register cells even when they were not active during a recording session. Hence, imaging a static neuronal marker along with dynamic GCaMP signal is a prominent application of a dual-channel Miniscope and enables researchers to study hippocampal spatial representation stability in an unprecedented way.

Here, we present a novel Miniscope that is capable of dual-channel calcium imaging in behaving animals. Our new design was derived from version 4 of the UCLA Miniscope with two high-sensitivity PYTHON480 CMOS sensors, 1mm x 1mm field of view, and electrowetting focus adjustment. The resolution of our dual-channel Miniscope is 3.5µm and all optics within the scope are achromatic (same as Miniscope v4). Our new design includes a sliding mechanism to correct any displacement in the focal plane across the two channels caused by chromatic aberration when using a GRIN lens to image deeper brain regions. The fully assembled dual-channel Miniscope weighs 4.8g and can be carried by freely behaving mice. To validate imaging of two wavelengths of light in freely behaving animals, and to demonstrate the application of the dual-channel Miniscope in the study of hippocampal spatial representations, we performed an imaging experiment where animals ran along a linear track across days. We injected a virus which co-expressed GCaMP and a constitutively active tdTomato in the CA1 region of the hippocampus. The constitutively active tdTomato signals provided a stable marker for cells that were within the imaging field of view. Our results are consistent with prior reports, suggesting a change in the hippocampal spatial representation over time even when the instability of field of view is accounted for using the additional tdTomato channel, demonstrating a prominent application of our dual-channel Miniscope.

## Results

### Dual-channel Miniscope design

We built our proof-of-concept dual-channel Miniscope to image GCaMP and tdTomato, but it could be used to image any two fluorophores by adjusting the appropriate filters and LED. Figure 1 shows a schematic of the design. To achieve simultaneous dual-channel imaging, we used a dichroic mirror to split the two emission wavelengths and used two CMOS sensors to acquire the images from two channels simultaneously. The excitation LED provides the light source for excitation. Depending on the specific application, different types of excitation LED can be installed that can either emit white (RGB), dual wavelengths, or even single wavelength excitation light, as long as it provides enough power to excite both fluorophores. Here, we used an LED with a single cyan wavelength that is strong enough to excite both GCaMP and tdTomato. The excitation light then travels through a dual-band-pass excitation filter that selectively passes through the two excitation wavelengths, followed by a half-ball lens to provide filtered and uniform excitation light. Next, the dual-band excitation dichroic designed to specifically reflect the two excitation wavelengths reflects the excitation light toward the sample. The excitation light then passes through the electrowetting lens followed by an objective achromatic lens stack and is focused onto the sample. The emission light (containing two wavelengths) travels through the achromatic lens stack and becomes collimated. By passing the excitation and emission lights through the same electrowetting lens and achromatic stack, we ensure that the focal plane is the same for excitation and emission and can be adjusted together using the electrowetting lens. The emission light then travels through the electrowetting lens and the dual-band excitation dichroic and is re-focused by the emission achromatic lens. The two wavelengths in the emission light are split by a high-pass emission dichroic. The higher wavelength passes through the emission dichroic and is collected by the CMOS sensor on the top, while the lower wavelength is reflected by the emission dichroic and collected by the CMOS sensor on the side. An emission filter is placed before each CMOS sensor to pass through the specific emission wavelength and minimize crosstalk between the two channels. Since all of the optics in the dual-channel Miniscope are achromatic, we placed the CMOS sensor in a way so that the distances traveled by both emission wavelengths are equal from the emission achromatic lens to the CMOS sensor, and the distance matches the focal length of the emission achromatic lens. This ensures that the focal plane is the same across both channels. Since the imaging principle is the same across different fluorophores with different excitation and emission wavelengths, our design is versatile to image a range of different combinations of fluorophores. The only modification needed to image a different set of fluorophores is to replace the excitation LED, excitation/emission filters, and dichroics to match the specific excitation/emission wavelengths of the fluorophores. Table S1 summarizes the bill-of-materials needed to assemble the dual-channel Miniscope, with the LED and filter set designed to image dynamic GCaMP signals and static tdTomato signals simultaneously. This configuration was used in the benchmarking and *in vivo* validation experiments described below.

**Figure 1:**
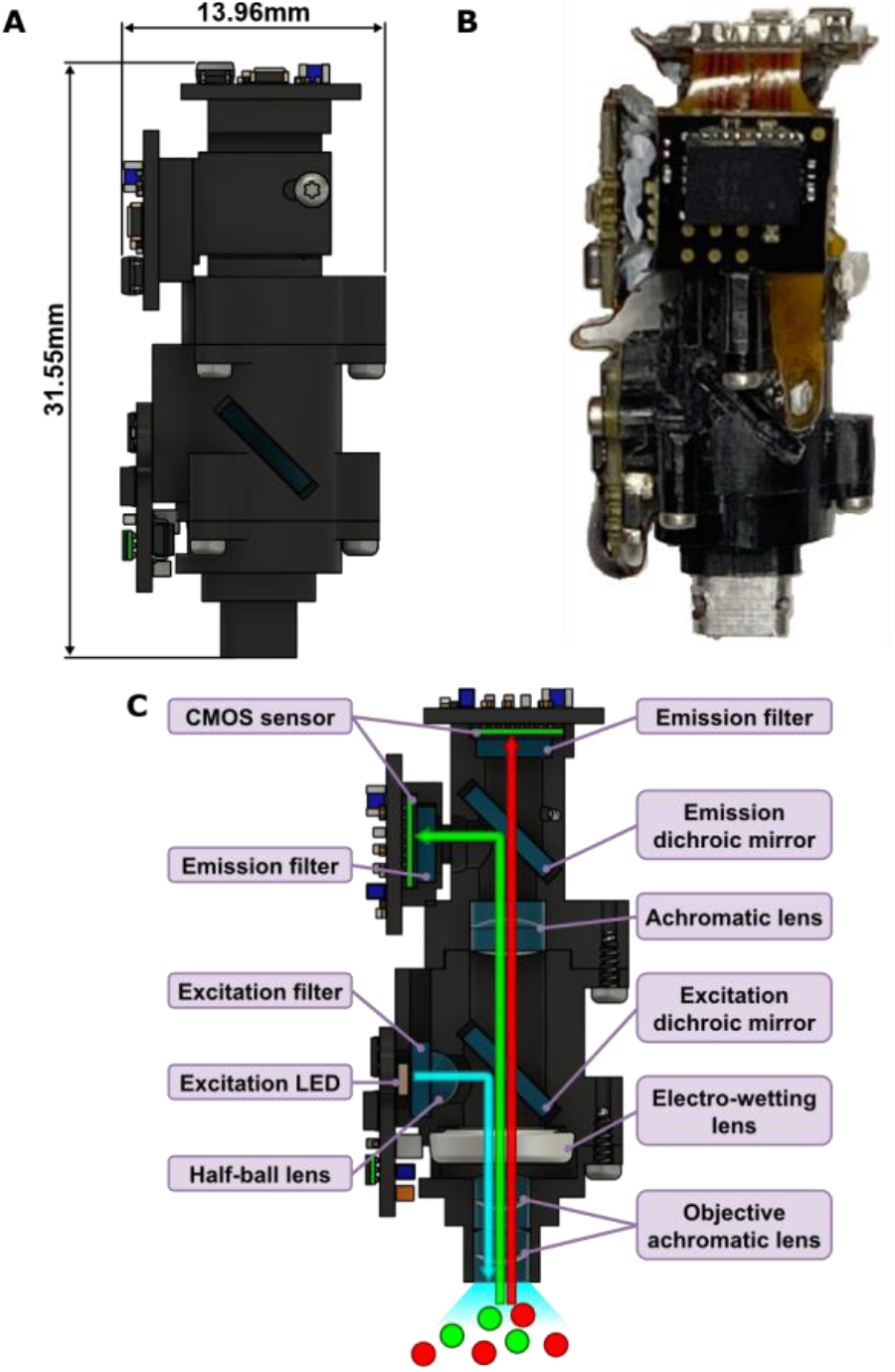
Dual-channel Miniscope design. **(A)** 3D rendering of the dual-channel Miniscope from the side showing the height (31.55mm) and diameter (13.96mm) of the Miniscope body. **(B)**Photo of the dual-channel Miniscope from the side. **(C)** Cross section of the dual-channel Miniscope showing the components and light path. The LED provides excitation light (cyan arrow). The excitation light travels through excitation filter and half-ball lens, then gets deflected by the dual-band excitation dichroic mirror, followed by the electro-wetting lens and objective achromatic lens stack. Once the sample is illuminated, the emission light with two wavelengths (green and red arrows) travels straight up into the dual-channel Miniscope. The two wavelengths pass through the objective achromatic stack, the electro-wetting lens, the excitation dichroic mirror, and are focused by the achromatic lens. The wavelengths are then split by the high-pass emission dichroic mirror, which deflects the lower wavelength (green) and passes through the higher wavelength (red). Finally, emission light is collected independently with two CMOS sensors with corresponding single-band-pass emission filters.

Although the dual-channel Miniscope is designed to be achromatic, in practice, a GRIN lens is often needed to relay and reproduce images from deeper brain regions to the working distance of the dual-channel Miniscope, which introduces chromatic aberration and causes a relative displacement in the focal plane across the two channels. Because the actual displacement depends on the pitch and diameter of the GRIN lens used, it is important to be able to correct for this displacement after assembling the dual-channel Miniscope. The design described here includes a sliding mechanism that allows the user to calibrate the focal plane of the two channels independently during a calibration imaging session (Figure 2). The CMOS sensor on the side is attached to a modular piece that fits onto the main body of the scope (Figure 2A). This modular piece is able to slide along the light path, changing the placement of the side CMOS sensor, which essentially changes the relative displacement between the focal planes across the two channels. The modular piece can be secured by two screws once the desired placement is determined, which remains stable throughout an imaging experiment. Using this mechanism, users can calibrate the focal planes of the two channels on the bench by imaging a test target through the specific GRIN lens used in the experiment. After calibration, we observed a high degree of overlap between the two channels both with and without a GRIN lens (Figure 2B), which suggests that this calibration process ensures imaging of the same focal plane even when imaging through a GRIN lens (Figure 2C).

**Figure 2:**
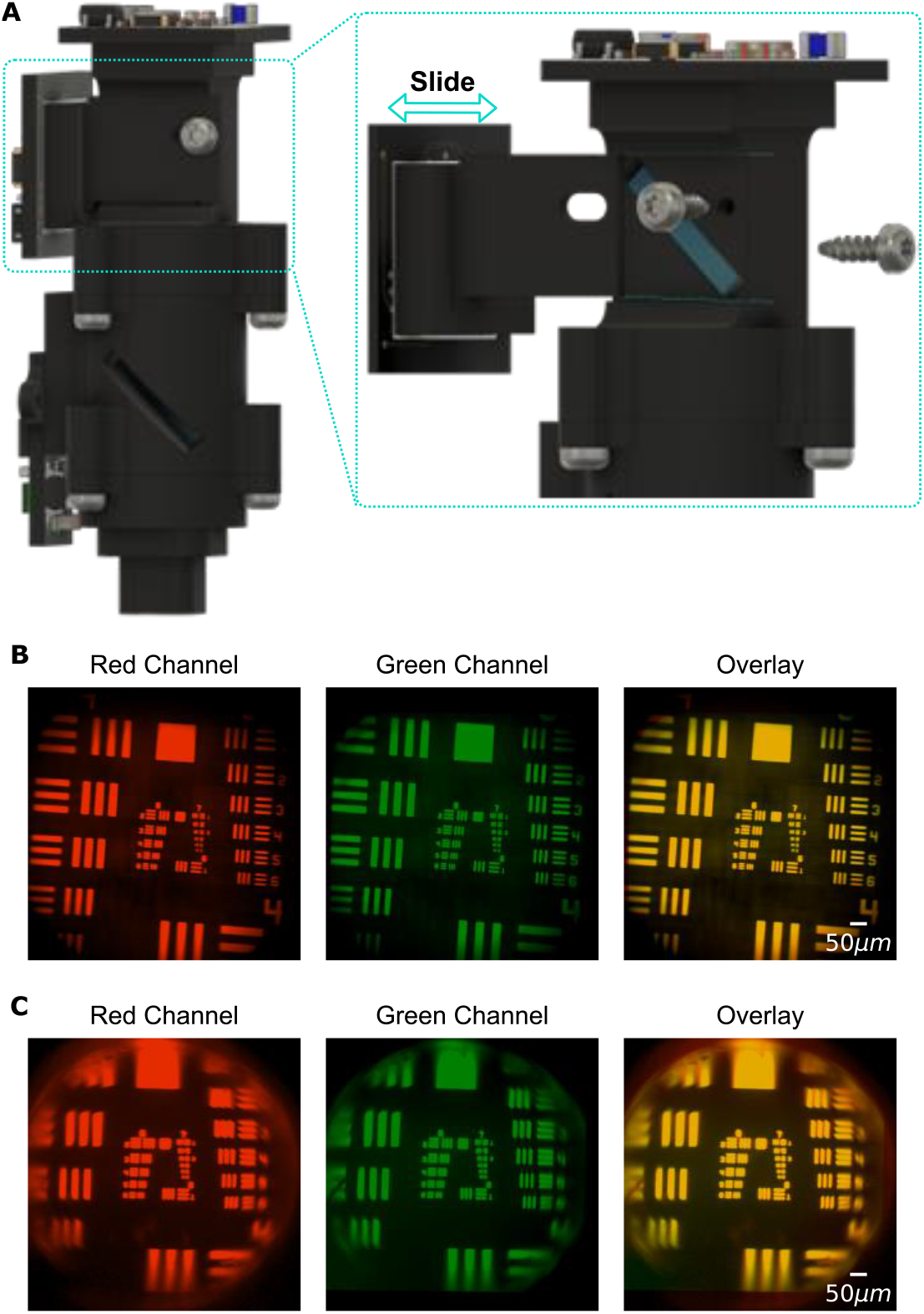
Sliding mechanism of the dual-channel Miniscope. **(A)** 3D rendering of the dual-channel Miniscope. The inset shows how the side CMOS sensor can slide along the optical path and be secured with two screws, which allows calibration of focal plane after assembly. **(B)** Image of a 1951 USAF resolution test target collected from red channel (top sensor) and green channel (side sensor) without a GRIN lens. An overlay of the two images demonstrates the same field of view across the two channels. **(C)** As in **B**, with the addition of a 0.5 pitch 1mm diameter GRIN lens between the scope and the test target. Deformation caused by the GRIN lens is visible near the edges of the images. The field of view is the same across the two channels despite chromatic aberration introduced by the GRIN lens.

### Validation of dual-channel Miniscope in freely behaving animals

To track the same population of cells across long periods of time, and to validate that we are able to image the same focal plane across two channels in freely behaving animals, we injected a virus that co-expressed GCaMP6f together with a static tdTomato in the same cell (Figure S1). We trained water-restricted mice to run on a linear track to retrieve 15μl of water rewards at both ends of the track while imaging hippocampal CA1 region using our dual-channel Miniscope (Figure 3A). Mice performed well on the linear track while wearing the dual-channel Miniscope (Figure 3B), reaching an average speed of 57.70 cm/s (± 2.22 cm/s) when running and ran an average of 41.42 (± 3.37) trials during 15 minutes of recording. To investigate the behavioral impact of the dual-channel Miniscope, we compared the running speed of animals (defined as 95^th^ percentile of locomotion speed when animals were not at reward zones, see [Star Methods]) in the dual-channel imaging experiment to another group of animals running on a linear track with a single-channel Miniscope. The average running speed of animals wearing dual-channel and single-channel Miniscope was 57.70 and 57.18 cm/s respectively, with no significant difference between the two groups (*p* = 0.873, independent t-test). Notably, to reach a similar behavioral performance with the dual-channel Miniscope, animals need to be habituated for longer (3 weeks instead of 1-2 weeks for single-channel Miniscope). This is likely due to the increased weight of the dual-channel Miniscope. Nevertheless, this result suggests that given enough training time, the dual-channel Miniscope has minimal behavioral impact for imaging experiments, similar to typical single-channel Miniscopes.

**Figure 3:**
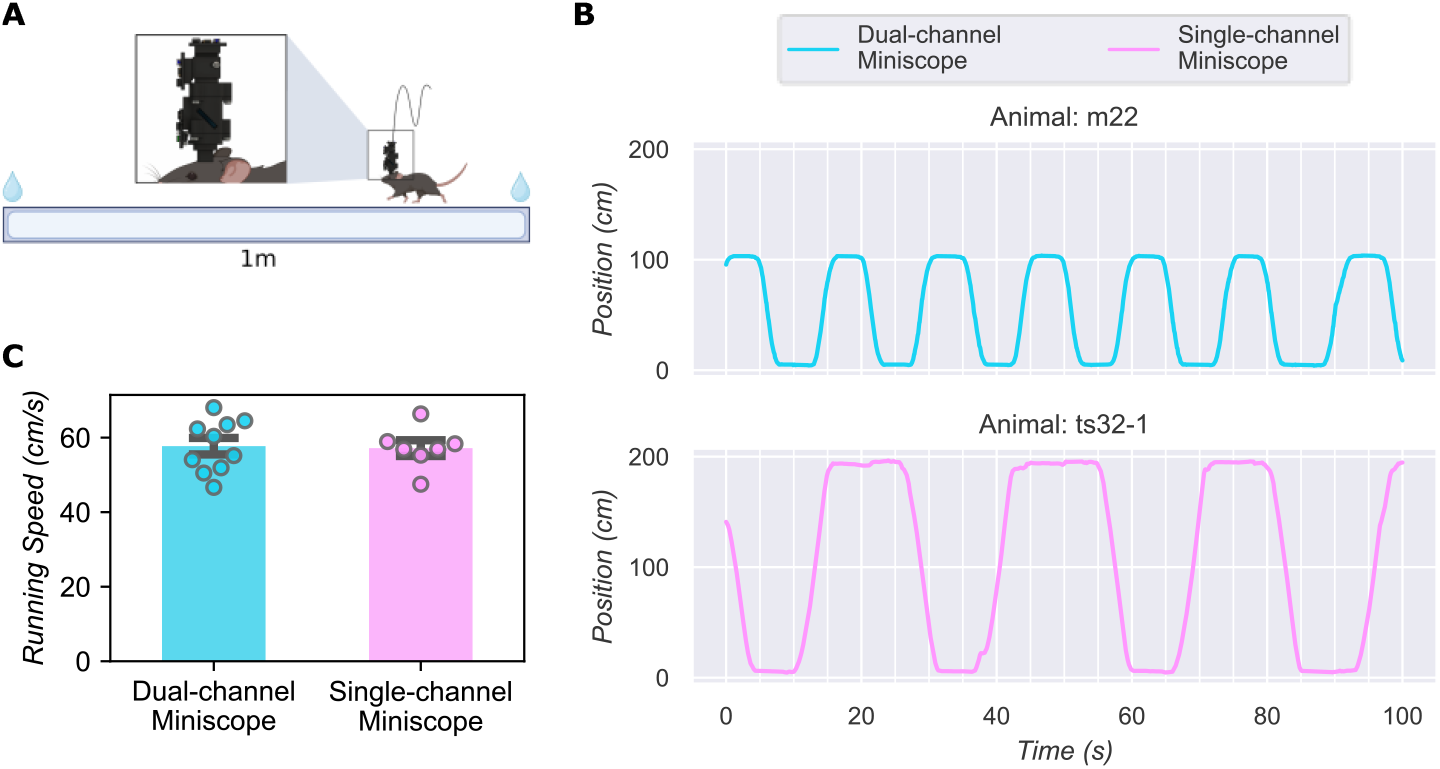
Imaging experiment using dual-channel Miniscope. **(A)** Schematic of the imaging experiment. Mice run on a 1-meter-long linear track to retrieve water rewards at both ends while wearing the dual-channel Miniscope. **(B)** Locomotion of representative animal wearing dual-channel or single-channel Miniscope while running on a linear track. For dual-channel Miniscope, the animals ran on a 1-meter-long linear track. For single-channel Miniscope, the animals ran on a 2-meter-long linear track. **(C)** Comparison of running speed between animals wearing single-channel Miniscope and animals wearing dual-channel Miniscope. The running speed was not significantly different between the two groups of animals.

To validate that we can track the same population of cells across a long period of time, we repeated the 15 minute-long recording with 2-day intervals between sessions for a total of 7 sessions, spanning 13 days. We then processed the GCaMP channel and tdTomato channel independently to extract the spatial footprints of cells. Next, we extracted the temporal activity of the detected cells. For the GCaMP channel, we applied the CNMF algorithm to extract the calcium dynamics and refined the spatial footprints of each cell. For the tdTomato channel, we projected raw fluorescence pixel values onto the spatial footprint of each cell. We then registered the footprint of each cell in the GCaMP channel to a footprint in the tdTomato channel based on the distance between centroids. An example field of view is shown in Figure 4A. The distribution and shape of spatial footprints were highly similar across the two channels (Figure 4A), suggesting that both channels are imaging the same field of view.

**Figure 4:**
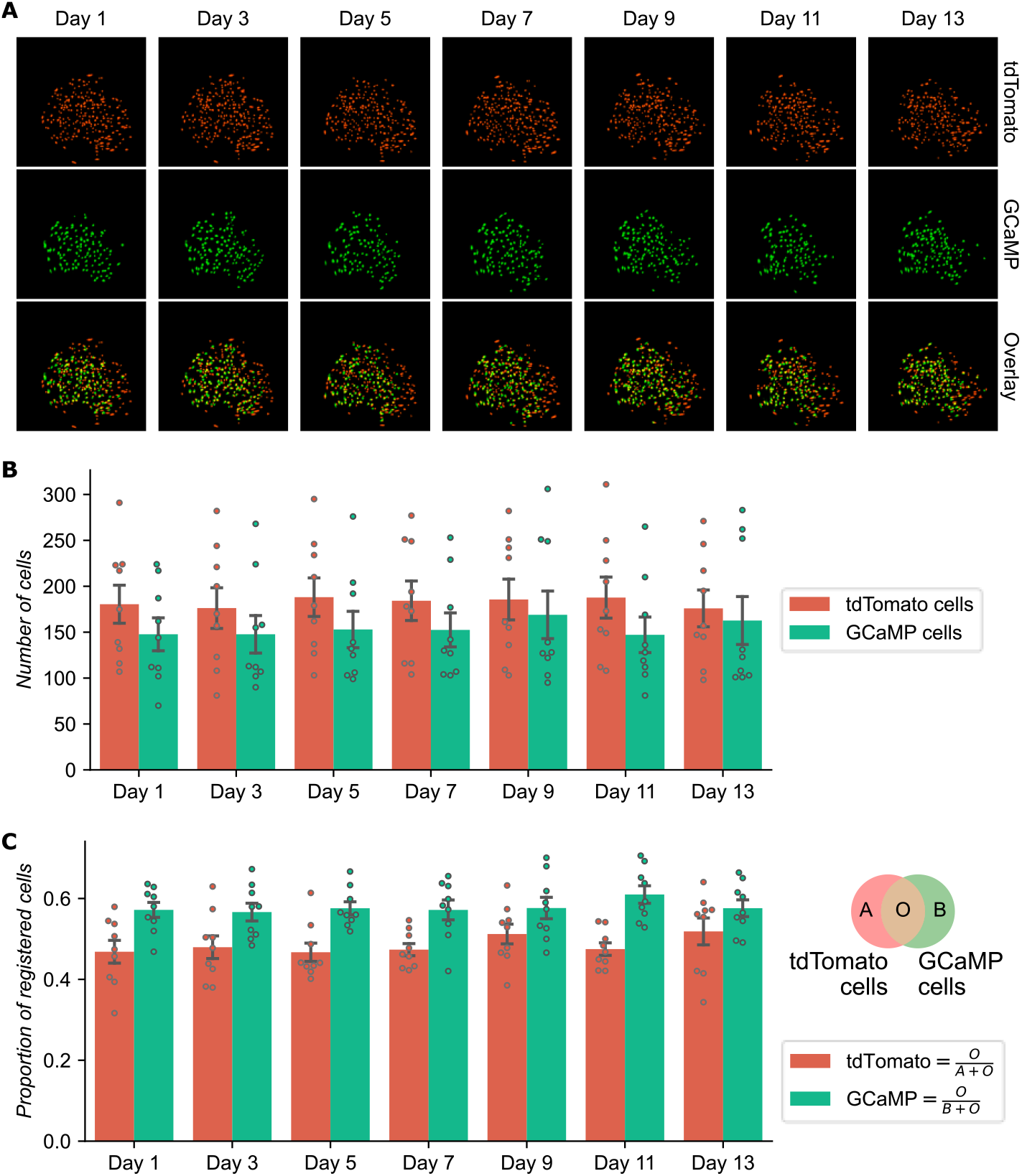
Cell registration across two channels. **(A)** Representative field of view from a single animal throughout the experiment. Processed spatial footprints are pseudo-colored and shown for the tdTomato (top) and GCaMP (middle) channels and the overlay images (bottom). Cells from the GCaMP channel that cannot be registered to the tdTomato channel are excluded from the plot. **(B)** Number of cells detected in each session for the tdTomato and GCaMP channels. **(C)**Proportion of cells registered across two channels. The venn diagram illustrates how the proportion was calculated for tdTomato and GCaMP channel in each session. The numerator was the number of registered cells across the two channels. The denominator was the total number of tdTomato and GCaMP cells for the tdTomato and GCaMP channel respectively.

We then quantified the number of cells detected in each recording session, as well as the number of cells that were registered across the two channels, as a fraction of total number of cells in either the GCaMP or tdTomato channel (Figure 4B, C). Each day, around 150 cells were detected in the field of view for each channel. Among these cells, approximately 50% of them can be registered, either as a fraction of GCaMP channel cell counts (Figure 4C, green bars) or tdTomato cell counts (Figure 4C, red bars). It is expected that not all tdTomato cells can be registered to a GCaMP cell in each day since hippocampal neural activity is sparse. However, in an ideal case, all GCaMP cells should have a corresponding cell in the tdTomato channel.

Several factors might explain why the proportion of registered cells did not reach 100% for the GCaMP channel. First, we are using an excitation LED that is optimized for GCaMP but not tdTomato (Table S1), so it is possible that not all cells were visible in the tdTomato channel each day. Second, different cells might have different relative expression levels for GCaMP and tdTomato, which could result in cells that were visible in the GCaMP channel but not tdTomato channel. Lastly, there is no temporal dynamic in the tdTomato channel, so we cannot use temporal dynamics to distinguish cells from background fluorescence and noise in the tdTomato channel. This makes it more challenging to detect cells in the tdTomato channel compared to the GCaMP channel. Importantly, none of these factors affect our ability to interpret neural activity from GCaMP cells that can be registered with the tdTomato channel.

To assess for crosstalk between channels, we investigated the cells registered across both channels and compared the corresponding temporal activity between the GCaMP and tdTomato channels. There was no visible change in the fluorescence in the tdTomato channel when a calcium event was visible in the GCaMP channel, despite the almost overlapping spatial footprints from the two channels, suggesting that there is minimal crosstalk from the GCaMP channel to the tdTomato channel (Figure 5).

**Figure 5:**
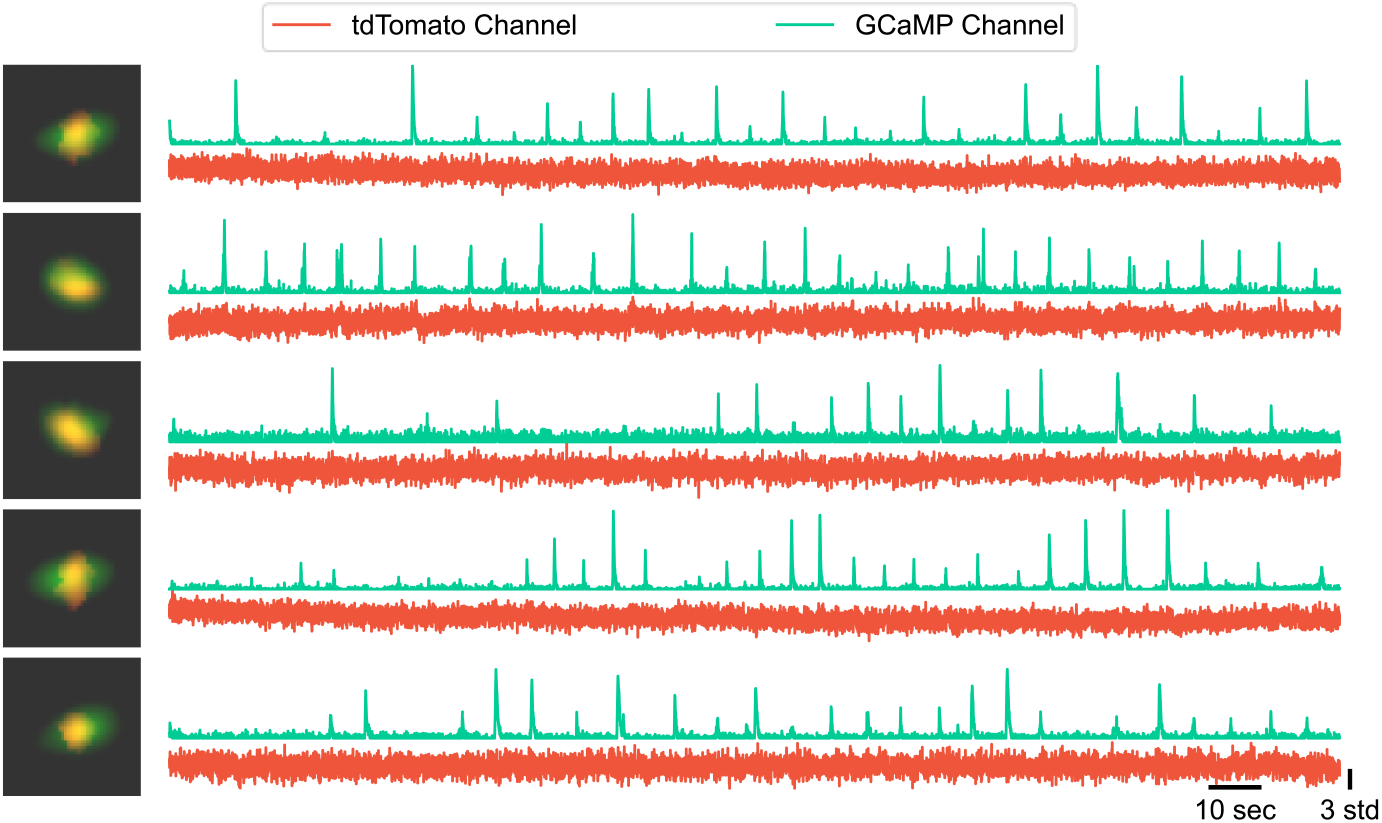
Representative cells registered across tdTomato and GCaMP channels. The processed spatial footprint for each cell is pseudo-colored for the tdTomato (red) and GCaMP (green) channels. A zoomed-in overlay image of the spatial footprint is shown for each cell on the left. The corresponding z-scored raw fluorescence from both channels is shown for each cell on the right. The raw fluorescence traces, obtained by projecting raw pixel values onto the highly overlapping spatial footprints, demonstrate minimal crosstalk between the channels.

To quantify crosstalk, we calculated an expected crosstalk ratio from the GCaMP channel into the tdTomato channel. The ratio is calculated as the amount of light in the GCaMP emission spectrum that passes through the tdTomato channel emission filter, divided by the amount of light passing through the GCaMP channel emission filter (Figure 6A). The resulting expected crosstalk ratio is 0.075 (i.e., in theory around 7.5% of fluorescence in GCaMP channel can be seen in the tdTomato channel). We then estimated the same crosstalk ratio from experimental data we collected. Leveraging the fact that the tdTomato channel should not contain any calcium dynamic in the ideal setup, we estimate the crosstalk ratio by linearly fitting the temporal dynamic of the tdTomato channel to that of the GCaMP channel for each cell registered across the two channels. Thus, the resulting linear model coefficient represented the amount of GCaMP dynamic that can be seen in the tdTomato channel as a fraction of those seen in the GCaMP channel, providing an estimation of the observed crosstalk ratio. The distribution of estimated crosstalk ratio is shown in Figure 6B. In addition, we generated a null distribution of the crosstalk ratio by randomly shuffling the temporal dynamic of one channel relative to the other for each cell registered across the two channels. The shuffled distribution centered around 0, representing no crosstalk (Figure 6B). The mean of the observed crosstalk ratio was significantly different from both the mean of the shuffled distribution (*p* < 0.001, paired t-test) and the expected crosstalk ratio (*p* < 0.001, one-sample t-test). These results suggest that although there is a non-zero amount of crosstalk, it is relatively minimal and less than the theoretical crosstalk ratio of 7.5%.

**Figure 6:**
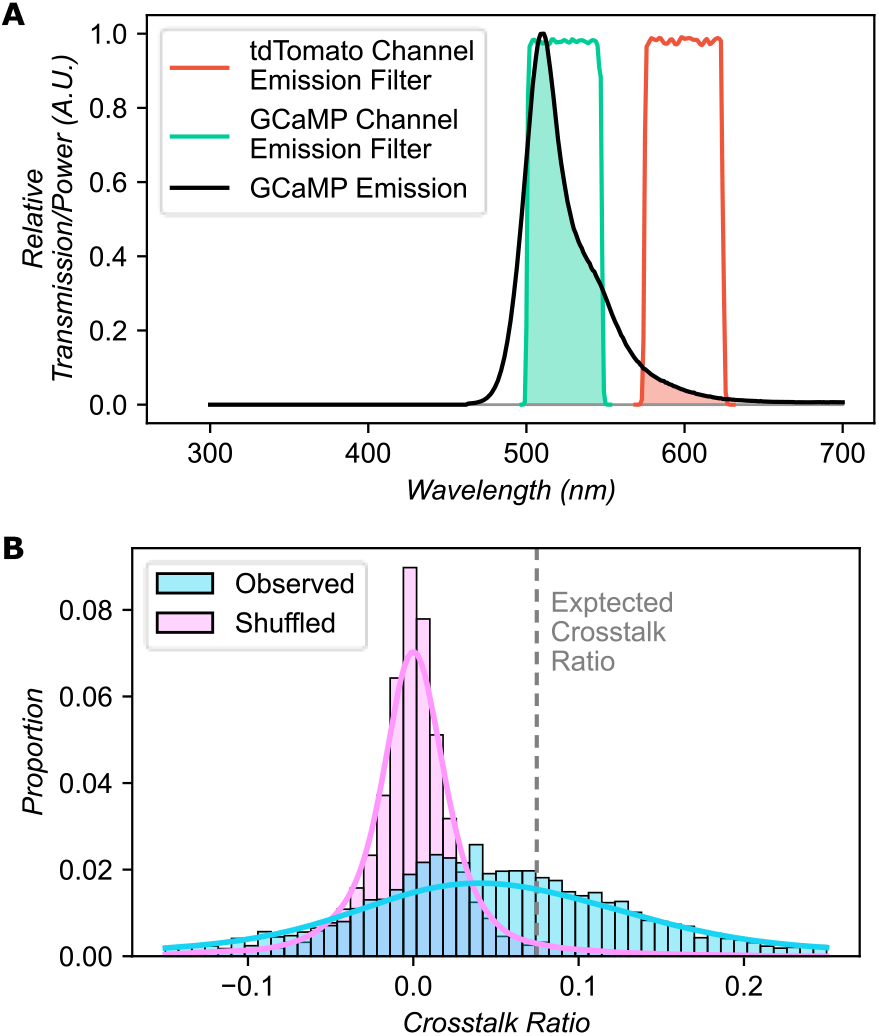
Quantification of crosstalk between the two channels. **(A)** Illustration of how the expected crosstalk ratio is calculated. The emission spectrum of GCaMP is shown in arbitrary units, overlaid with the relative transmission profile for the GCaMP and tdTomato channel emission filter. Green shaded area indicates the relative amount of GCaMP emission that passes through the GCaMP channel filter; red shaded area indicates emission passing through the tdTomato channel filter. The expected crosstalk ratio is calculated as the ratio between the red and green areas. **(B)** Distribution of crosstalk ratio estimated from experimental data. The distribution of estimated crosstalk ratio (linear model coefficient) is shown in cyan, with the shuffled distribution overlaid in purple. The expected crosstalk ratio is also shown as dashed line.

Together, these results suggest that using our dual-channel Miniscope, we can image the same field of view across two channels with minimal crosstalk in freely behaving mice. Taking advantage of an additional static channel, we can track large population of cells in the hippocampal CA1 region across weeks.

### Investigation of hippocampal neural ensemble stability using the dual-channel Miniscope

With the capability to track a large population of CA1 cells across weeks to months, we further investigated the stability of hippocampal spatial representations. The change in hippocampal spatial representations has been reported in two distinct ways. First, the reactivation rate of the neural population decreases as the time interval increases, suggesting that different populations of cells are active at different times [4,27]. Second, the spatial population vector correlation decreases as the time interval increases, suggesting that the spatial representation is constantly changing over time [4,27]. Both metrics can be influenced by instability of the field of view, potentially resulting in over-estimation of the rate of changes over time. Hence, the instability of the field of view must be accounted for when studying the long-term stability of hippocampal spatial representations.

To account for instability in the field of view, we used the additional static tdTomato channel and studied the subset of cells that we can stably track across all 13 days of the imaging experiment. We investigated the stability of cells in each channel by quantifying (on a per-session basis) the probability of a cell being tracked in a different number of sessions (Figure 7). Overall, we were able to track a large proportion of cells through all 7 recording sessions in the tdTomato channel. In each session, on average 46.4% of the cells were stably tracked in the tdTomato channel throughout all 7 imaging sessions. There was a significant group (number of sessions) effect (*F*_6,56_ = 179.975, *p* < 0.001, type 3 ANOVA), and the per-session probability of cells tracked across 6 and 7 sessions were significantly different from the rest of probabilities (null hypothesis were rejected in all pairwise Tukey’s HSD tests involving given group). Notably, not all cells in the tdTomato channel can be tracked throughout all sessions, even though tdTomato expression is constitutive. This can be explained by the possibility that different cells might have different levels of tdTomato expression through time, or that not all cells in the field of view can be detected in the tdTomato channel due to previously mentioned reasons. An alternative explanation is that there might be instability in the field of view, where cells may move in and out of the field of view, driving down the probability of active cells across multiple sessions, highlighting the need for a constitutively active cell marker. As a comparison, when registering cells using only the GCaMP channel, the probability of cells active across all 7 sessions decreased to 22.2%. There was still a significant group (number of sessions) effect (*F*_6,56_ = 15.212, *p* < 0.001, type 3 ANOVA), and the per-session probability of cells tracked across 7 sessions were significantly different from the rest of probabilities (null hypothesis were rejected in all pairwise Tukey’s HSD tests involving a given group). It is not surprising that a smaller proportion of cells were active across all 7 recording sessions in the GCaMP channel. This is likely due to the sparsity of hippocampal neural activity, as not all cells are constantly active in any given session. However, it is still possible that instability of field of view may contribute to instability of neural activity across sessions. Because of this, we then focused our analysis on a subset of GCaMP cells, where the GCaMP cells can be registered to a stable subset of tdTomato cells that can be tracked across all 7 sessions. By extension, we call this subset of GCaMP cells the “stable GCaMP cells”, in the sense that we can be confident in their presence in the field of view, whether they were active or not, despite potential instability of field of view. Interestingly, in the stable GCaMP cells, on average only 19.7% of these cells were active across all 7 sessions, which is lower than the probability from all GCaMP cells. There was still a significant group (number of sessions) effect (*F*_6,56_ = 3.335, *p* = 0.007, type 3 ANOVA), but the probability of cells tracked across 7 sessions were not significantly different from all other probabilities (pairwise Tukey’s HSD test). Importantly, instability of field of view can no longer account for this result because the cells are all registered with tdTomato channel which ensures they are still present in the field of view across all sessions. Taken together, these results suggest that hippocampal neural activity was indeed sparse and different cells were active at different times even when instability of the field of view was accounted for.

**Figure 7:**
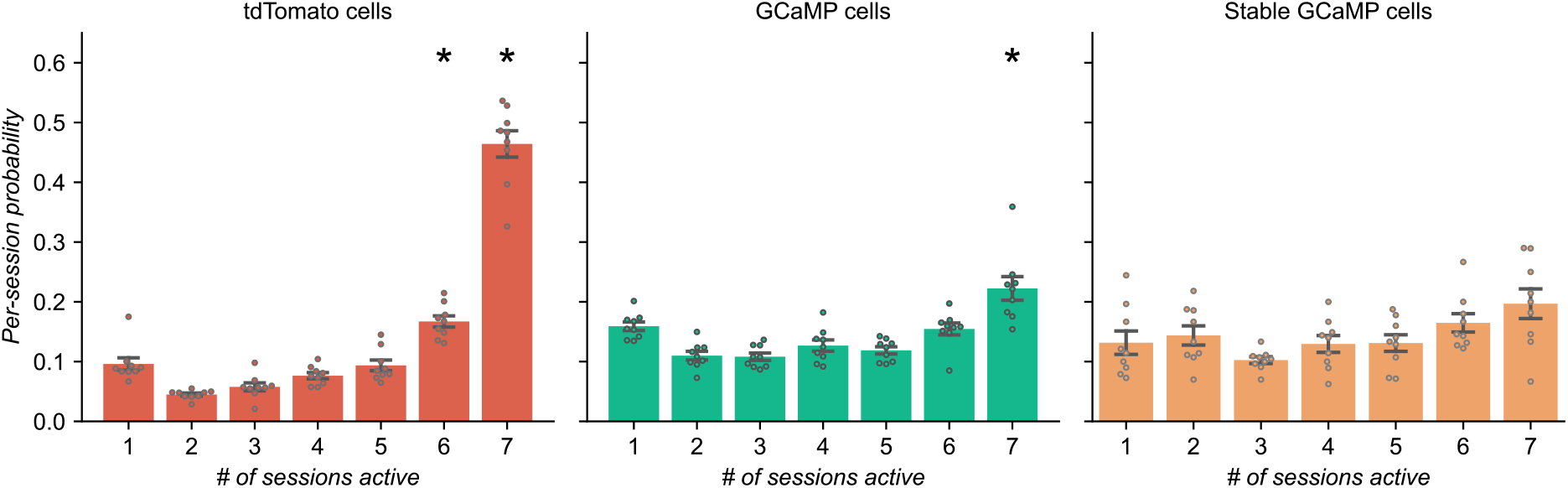
Tracking cells across weeks. Average per-session probabilities of cells tracked through different number of sessions are shown. The probability is estimated for each individual session as the number of cells tracked through certain number of sessions divided by the total number of cells in the session. The probability is then averaged across all sessions. The tracking result using the tdTomato channel is shown on the left. The tracking result using only the GCaMP channel is shown in the middle. Then, we registered each GCaMP cell to a corresponding landmark tdTomato cell, and aggregated only the GCaMP cells whose corresponding tdTomato landmark can be tracked stably across all 7 recording sessions. The tracking result of this stable subset of GCaMP cells is shown on the right. Star indicates a particular group (per-session probability for a particular number of sessions) was different from all other probabilities in post-hoc pairwise Tukey’s HSD tests.

Next, we focused our analysis on place cells [28,29] (see definition of place cells in [Star Methods]) and investigated the stability of the hippocampal neural ensembles as a function of time interval. As shown in Figure 8A, the reactivation probability of cells across pairs of sessions decreased as a function of time interval when we include all GCaMP cells (slope: -0.0217, *p* < 0.001, OLS). This result is consistent with previous findings on long-term stability of hippocampal spatial representation [4]. As a comparison, when we only included the stable GCaMP population, the reactivation rate was much higher compared to the average of all cells registered with the GCaMP channel. As expected, the reactivation rate still decreased as a function of time for cells registered with tdTomato but to a lesser degree (slope: -0.0127, *p* < 0.001, OLS). Overall, there were significant effects of time interval (*F*_1,109_ = 67.74, *p* < 0.001, type 3 ANOVA), cell inclusion (*F*_1,109_ = 5.72, *p* = 0.019, type 3 ANOVA), and the interaction between the two factors (*F*_1,109_ = 5.64, *p* < 0.019, type 3 ANOVA). We expected that reactivation probability would be lower when including all GCaMP cells, because including cells that cannot be stably tracked essentially increases the denominator during the calculation of probability, yielding a numerically smaller result. Taken together, these results suggest that, consistent with previous findings, the reactivation probability of cells decreases as a function of time interval, even when the instability of field of view is accounted for using static landmarks in the tdTomato channel.

**Figure 8:**
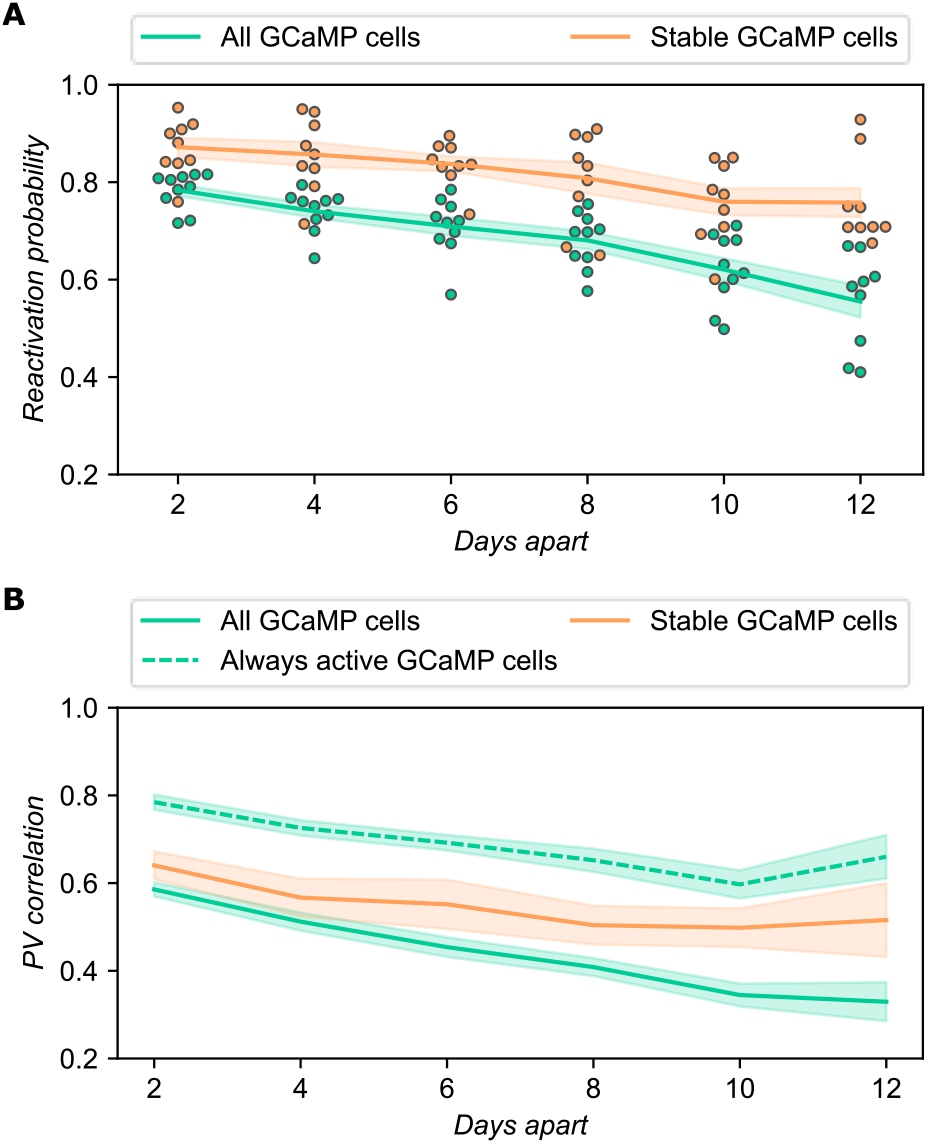
Hippocampal spatial representation changes over time. **(A)** Reactivation probability of cells across pairs of sessions as a function of time interval. The reactivation probability of cells for a pair of sessions is calculated as the number of cells shared across both sessions divided by the total number of cells in either session, then averaged across the pair of sessions. Solid lines show the mean across all possible combination of sessions with certain time interval and all animals, and shaded area shows the standard error of the mean. Green: all cells in GCaMP channel are included. Orange: only GCaMP cells whose corresponding tdTomato landmark can be tracked stably across all sessions are included. **(B)** Population vector correlation across pairs of sessions as a function of time interval. The population vector correlation is calculated as the mean Pearson correlation of population vectors across all spatial bins between a pair of sessions. Then a mean is taken within each animal for a given time interval across all possible combinations of recording session pairs. Lines show the mean across all animals and shaded area shows the standard error of the mean. This calculation is carried out across all GCaMP cells (green solid line) and cells whose corresponding tdTomato landmark can be tracked stably across all sessions (orange solid line). As a comparison, the same calculation is carried out with GCaMP cells that were active in both sessions in a pair regardless of the corresponding tdTomato landmarks (green dashed line).

After examining the reactivation of hippocampal neural ensembles, we proceeded to specifically investigate the stability of hippocampal spatial representation. As shown in Figure S2, within each day, the hippocampal place cells ensemble formed a spatial representation of the linear track, but the representation was not stable across days. This pattern was consistent regardless of whether we included all place cells found in the GCaMP channel or if we included only the place cells in the stable GCaMP population. We then calculated the spatial population vector correlation across days using place cells (Figure 8B) with different cell inclusion criteria based on the two channels. Either all GCaMP cells (Figure 8B, green solid line) or the stable GCaMP population (Figure 8B, orange solid line) were considered. As an additional comparison, we also carried out the same analysis for GCaMP cells that were active during both sessions in the pairwise calculation regardless of the corresponding tdTomato landmarks. As shown in Figure 8B, the population vector correlation decreased as the time interval increased for all three inclusion criteria (all GCaMP cells slope: -0.026, *p* < 0.001; stable GCaMP cells slope: - 0.013, *p* = 0.048; always active GCaMP cells slope: -0.015, *p* < 0.001, OLS). Overall, there was a significant effect of time interval (*F*_1,166_ = 51.31, *p* < 0.001, type 3 ANOVA) and inclusion criteria (*F*_2,166_ = 8.02, *p* < 0.001, type 3 ANOVA). However, when we excluded the “always active GCaMP cells” group, then there was no longer a significant effect between the two remaining inclusion criteria (*F*_1,109_ = 0.046, *p* = 0.830, type 3 ANOVA), while the effect of time interval remained significant (*F*_1,109_ = 33.09, *p* < 0.001, type 3 ANOVA). This suggests that the PV correlation was not significantly different regardless of whether we registered the GCaMP cells with tdTomato landmarks and included only the stable population. The numerically elevated PV correlations when including only always active GCaMP cells was again expected, since by including only active cells we essentially eliminated the zero terms introduced by inactive cells during correlation calculation. Consistent with previous findings, these results suggest that the hippocampal spatial representation changes over time, even when instability in field of view was accounted for. However, without registering GCaMP cells with the tdTomato channel, one cannot assume that cells not present in the field of view were inactive. Therefore, without the tdTomato channel, only the active cells expressing GCaMP can be included in the PV correlation analysis, which can result in an over-estimation of PV correlations.

In summary, we found that the hippocampal neural ensembles changed over time in terms of both the population of cells active and the spatial coding of the environment. This finding is consistent with previous reports and holds true even if we account for instability in imaging field of view using the tdTomato channel, although a more accurate rate of change can be obtained using the tdTomato channel.

## Discussion

### Dual-channel Miniscope

Here we presented a novel dual-channel Miniscope that can image two fluorophores in behaving animals. Our design includes a mechanism to calibrate the focal plane of the two channels and is lightweight enough to be carried by adult mice with minimal impact on behavior. We have validated our design at the bench and *in vivo* in freely behaving animals performing a spatial navigation task. We found that we can image the same field of view across days and the same focal plane across the two channels with minimal crosstalk, while the animals freely behave, wearing the dual-channel Miniscope. Furthermore, we demonstrated a use case where we take advantage of an additional static channel to track the same population of neurons even when some neurons were inactive in some recording sessions. We used the dual-channel Miniscope to show that the hippocampal spatial representation changes over time, consistent with previous findings. Importantly, we were able to account for instability in field of view, providing a more credible and accurate estimation of rate of change in the hippocampal neural ensemble.

### Comparison with alternative designs

Our dual-channel Miniscope uses two CMOS sensors to achieve simultaneous dual-channel imaging. Other work has used one CMOS sensor combined with dual-band-pass filters/dichroic and two excitation LEDs [30], where one of the LEDs turns on at every other frame and illuminates the sample at one of the two excitation wavelengths, resulting in a multiplexed imaging signal at half the frame rate. Compared to the single CMOS sensor design [30], our design allows us to separate the channels into two independent light paths and use two single-band-pass emission filters. Since all the filters used in Miniscopes are designed for collimated light, they often suffer from crosstalk due to imperfectly collimated light. The extent of crosstalk is often much more severe for dual-band-pass filters compared to single-band-pass ones. Hence, using two single-band-pass emission filters in our design provides better filtering quality and minimizes crosstalk between the two channels. The other benefit of having two CMOS sensors is that the relative positions of the two sensors can be easily adjusted to correct for any displacements in the focal plane potentially caused by chromatic aberrations. Further, full frame rates supported by the image sensors can be used since there is no need to multiplex the imaging data from the two sensors. The cons of using two CMOS sensors are increased physical dimensions and substantial increase in weight. However, we have validated our scope in behaving animals and our results suggest that after a period of training, mice can perform normally in a spatial navigation task.

### Limitations and future directions

Currently, our dual-channel Miniscope uses an excitation LED with a single peak wavelength, which was sufficient to excite both GCaMP and static tdTomato markers. Hence, to this end, we have only demonstrated one use case for our dual-channel Miniscope: using an additional static channel to image a constitutively active tdTomato fluorophore (together with GCaMP). This allowed us to account for the instability in imaging field of view and track the same population of neurons longitudinally, even when some neurons were not always active. However, our design can be modified for additional applications. For example, imaging a cell-type specific or activity-dependent marker together with calcium activity can reveal the different activity patterns for neurons with different functions. Furthermore, having two channels opens up the possibility of fluorescence resonance energy transfer (FRET) microscopy using a Miniscope. Cell-based neurotransmitter fluorescent engineered reporters (CNiFERs) that emit FRET signals have recently been developed to detect the release of neurotransmitters [31,32,33]. Our dual-channel Miniscope can be modified to image numerous FRET-based CNiFERs to study the dynamics of neurotransmitters. Finally, a researcher might image two dynamic fluorescence signals across the two channels, e.g., using GCaMP and RCaMP (a red-shifted calcium indicator) to mark excitatory and inhibitory neurons, respectively, to study the distinct population dynamics. One can also image GCaMP signals from neurons together with dynamics of neurotransmitter release using novel genetically encoded sensors [34,35,36,37]. To image these different combinations of fluorophores, usually either a dual-wavelength LED or two different LEDs are required. In both cases, two independent LED drivers need to be integrated on the PCB, which is feasible since there is extra space on the PCB in our current prototype and the power drawn from additional LEDs is relatively small. Hence, the next step in future development of our dual-channel Miniscope is to enable two excitation wavelengths, which greatly expand the applicability of our dual-channel Miniscope.

The dual-channel Miniscope weighs 4.8 grams, which can be carried by mice but requires a much longer period of training time comparing to single-channel Miniscopes. The fact that two coax cables are required for the current dual-channel Miniscope prohibits the use of typical commutators, which exacerbates the tether of animals and contributes to the longer training time. To decrease the overall weight of our dual-channel scope and provide compatibility with typical commutators, the PCBs could be designed so that the image sensors and other components are integrated more compactly on a single PCB. Currently, the PCBs used in our dual-channel Miniscope are two independent PCBs from Miniscope v4 intended to serve as a proof-of-concept, which means that many ICs are duplicated and add unnecessary weight to the Miniscope. In particular, two serializers on the two separate PCBs necessitates two separate coax cables. Future iterations could include a one-piece PCB that hosts two excitation LEDs and two image sensors, with only one copy of the rest of the components including the electro-wetting lens driver, microcontroller and serializer. Hence, future development of the dual-channel Miniscope can further reduce the weight and provide more flexibility in experimental setups.

To date, we have only validated the dual-channel Miniscope with a limited number of experiments. More validations are needed to ensure our dual-channel Miniscope is suitable for broader applications. For example, to fully validate that we can image the same focal plane in behaving animals, we need to express two static markers with different emission wavelengths in the same population of neurons. A high degree of overlap in this case would indicate successful imaging of the same population of neurons across two channels, although this too could be subject to other confounds such as expression-level differences as mentioned. Furthermore, we could image two dynamic calcium indicators (for example, GCaMP and a red-shifted calcium indicator such as RCaMP or jRGECO) in the same field of view with our dual-channel Miniscope. We could image the two calcium indicators in the same population of neurons and calculate the correlation of temporal dynamics of the matching cells across the two channels. A high degree of correlation would further confirm that we can image the same focal plane and the signals are not contaminated by out-of-focus fluorescence, provided the effect of crosstalk can be ruled out. Additionally, we could image two calcium indicators in different, non-overlapping subsets of neurons and again investigate the correlation between temporal signals at matching locations in the field of view across two channels to evaluate crosstalk. Based on our validation of limited crosstalk between the two channels, we expect no significant correlation of the temporal signals. Still, this is an important validation to confirm that we can image two dynamic signals simultaneously with minimal crosstalk.

In conclusion, we have demonstrated that our dual-channel Miniscope is capable of imaging two fluorescence wavelengths in behaving animals. We believe our dual-channel Miniscope has many applications and a significant impact on modern neuroscience research.

## Materials and Methods

### Surgery and imaging experiment

All experimental protocols were approved by the Institutional Animal Care and Use Committee of the Icahn School of Medicine at Mount Sinai in accordance with the US National Institutes of Health (NIH) guidelines. 10 adult male C57BL/6J mice were used during the validation imaging experiment. AAV1-hSyn-GCaMP6f-P2A-nls-dTomato virus (titer 2 × 10^13^ GC/mL) that co-expresses GCaMP and nucleus-localized static tdTomato signals was purchased from Addgene, which was a gift from Jonathan Ting (Addgene viral prep #51085-AAV1; http://n2t.net/addgene:51085; RRID:Addgene_51085). For all surgeries, mice were anesthetized with 1.2-1.5% isoflurane and head fixed onto a stereotax (David Kopf Instruments). Lidocaine (2%, Akorn) was injected to the neck area as an analgesic. All mice underwent virus injection and lens implantation stereotaxic surgeries on the same day. First, mice were unilaterally injected with 500 nl of AAV1-hSyn-GCaMP6f-P2A-nls-dTomato virus at 2nl/sec in the dorsal CA1 (−2.0 mm anteroposterior relative to bregma, +1.8 mm mediolateral from bregma, and 1.6 mm ventral from the skull surface) using a Nanoject microinjector (Drummond Scientific). Next, mice underwent a GRIN lens implantation surgery. A craniotomy at 1 mm in diameter was performed above the viral injection site. The cortical tissue above the corpus callosum was carefully aspirated using 27 gauge and 30 gauge blunt needles while buffered artificial cerebrospinal fluid (ACSF: in mM: 135 NaCl, 5 KCl, 5 HEPES, 2.4 CaCl_2_, 2.1 MgCl_2_, pH 7.4) was constantly applied throughout the aspiration to prevent desiccation of the tissue. The aspiration ceased after partial removal of the corpus callosum and full termination of bleeding, at which point a GRIN lens (1 mm diameter, 4.0 mm length, 0.5 pitch, Inscopix) was stereotaxically lowered to the targeted implant site (−1.35 mm dorsoventral from the bregma). Cyanoacrylate glue and dental cement was used to seal and cover the exposed skull, and Kwik-Sil (World Precision Instruments, CAT. 600022) covered the exposed GRIN lens. A final layer of cement was then applied above the Kwik-Sil to protect the lens from group housing. Subcutaneous saline injections were administered at the end of each surgery to prevent dehydration. Carprofen (5mg/kg), dexamethasone (0.2 mg/kg), and ampicillin (20mg/kg) were administered during surgery and for 7 days after surgery together. Animals were anesthetized again 3 weeks later, and the dual-channel Miniscope with an aluminum baseplate attached was placed on top of the GRIN lens. After searching the field of view for in-focus cells under the tdTomato channel, the baseplate was cemented into place and the Miniscope was unlocked and detached from the baseplate. A plastic cap was locked into the baseplate to prevent debris build-up.

During the imaging experiment, mice were water-restricted by replacing the water supply with 4mg/ml citric acid [38]. All mice maintained 80% - 85% of their original body weights throughout the imaging experiment. Mice were habituated to run on a 1-meter linear track to retrieve water rewards at both ends for 2 weeks before the imaging experiment. Each visit to the two ends of the track was considered one trial. Mice wore custom-made dummy Miniscopes with increasing size and weights through the course of habituation until all mice could run at least 10 trials on the linear track within 15 minutes wearing the dual-channel Miniscope with cable attached. The imaging experiment consisted of seven 15 minutes imaging sessions with a two-day interval, spanning a total of 13 days. The behavior video was collected with an off-the-shelf webcam (ELP-USBFHD01M-L180, Amazon). The imaging data were collected with standard Miniscope v4 data acquisition hardware and software (https://github.com/Aharoni-Lab/Miniscope-DAQ-QT-Software).

To compare the running speed of mice wearing the dual-channel Miniscope with mice wearing a traditional v4 Miniscope, we collected data from another group of mice that underwent the same surgical and experimental procedures. This group of mice was trained to collect water rewards on a 2-meter-long linear track while wearing single-channel Miniscope v3. To estimate the animals’ running speed, we calculated the gradient of position change (in pixels) for each frame, then scaled the value to physical units (cm/s) based on the track length and video frame rate. We excluded all frames when the animals were in the reward zones and used the 95^th^ percentile running speed of the rest of the frames as the estimated running speed for a session to avoid biasing results with acceleration or deceleration.

### Data analysis

Animal positions were tracked with ezTrack [39,40] (https://github.com/denisecailab/ezTrack) and the calcium activity from the GCaMP channel was processed using Minian [41] (https://github.com/denisecailab/minian). The static tdTomato channel was processed using a modified pipeline based on Minian, which detected cells by locating local maxima, then extracted spatial footprints by detecting round features around the local maxima. For the hippocampal population analysis (Figure 7), we excluded one animal (m30) because one recording session was missing for that animal. All downstream analyses were performed using in-house Python scripts that are available at https://github.com/denisecailab/Miniscope_2s-validation. Briefly, after extracting the spatial footprints of cells from both tdTomato and GCaMP channels, we extracted denoised calcium activity and deconvolved signal using Minian from the GCaMP channel. To benchmark the crosstalk between the two channels, we extracted the equivalent of temporal activity for each cell from the tdTomato channel by projecting raw pixel fluorescence values onto the spatial footprints. Next, we registered cells across recording sessions using a centroid-distance based registration algorithm in Minian. We registered cells using either the tdTomato channel or the GCaMP channel. Additionally, we registered each GCaMP cell to a tdTomato cell within each session. We then generated a cross-session registration mapping for GCaMP cells using the cross-session mapping of their corresponding tdTomato cells. In this way, the tdTomato cells serve as stable landmarks across recording sessions even when the cell was not active and not detected in the GCaMP channel in particular sessions.

To calculate the per-session probability of cells tracked through different sessions, we used the equation

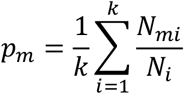

where *p*_*m*_ is the proportion of cells active across *m* recording sessions, *k* is the total number of recording sessions (i.e. *k* = 7 when looking across the whole experiment), *N*_*i*_ is the total number of cells detected in session *i*, and *N*_*mi*_ is the number of cells in session *i* that are also tracked across exactly *m* sessions. In other words, this calculation estimates the probability of finding a cell being tracked through a different number of sessions on a per-session basis. Similarly, we can derive the reactivation rate as a special case where we were only looking at a pair of sessions and we calculated probability of finding a cell that is active in both sessions in either session, in other words, *m* = 2 and *k* = 2. Expanding the summation and explicitly labeling the two sessions as *A* and *B*, we have:

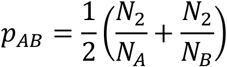

where *p*_*AB*_ is the reactivation rate regarding session *A* and *B, N*_*A*_ and *N*_*B*_ are the total number of cells in session *A* and *B* respectively, and *N*_2_ is the number of cells active in both sessions.

To investigate the hippocampal spatial map, we classified behavior episodes into idle, running left, and running right using a speed threshold of 3 cm/s. We then linearized the position of animals along the track and mapped the positions to the range of 0-100. To account for directionality of hippocampal place cells, we mapped the position independently when the animals were running left or right, resulting in a total range of 0-200. We used kernel density estimation to estimate both the spatial firing activity from the deconvolved signal and the occupancy from animal positions. A cosine kernel with a bandwidth of 5 cm was used for both spatial firing activity and occupancy. To estimate occupancy, the position of the animal at each frame when the animal was not idle was used to directly estimate the density of the positions (occupancy). To estimate spatial firing activity for each cell, the position of an animal at frames when there was a calcium event (i.e., non-zero value in deconvolved signal) was used and weighted according to the amplitude of the deconvolved signal to estimate the density of firing activity as a function of position. After kernel density estimation, we evenly sampled 200 spatial bins from range 0-200 (including both running directions), equivalent to a resolution of 1 cm/bin. We used the discretely sampled spatial firing activity and occupancy for all further calculation and plotting. To calculate spatial information, we used the equation

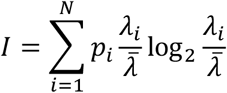

where *N* = 200 is the total number of spatial bins, *p*_*i*_ is the occupancy at spatial bin *i, λ*_*i*_ is the spatial firing activity at spatial bin *i*, and 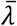 is the mean firing activity across the whole recording session. Next, we calculated normalized spatial firing activity by dividing the spatial firing activity at each spatial bin by the corresponding occupancy. Then, to calculate spatial stability within each recording session, we computed the mean Pearson correlation of normalized spatial firing activity across odd and even trials as well as first and second halves of trials. To calculate spatial stability across recording sessions, we computed the mean Pearson correlation of normalized spatial firing activity for the same registered cell across all combinations of sessions. To compute the activity level for each cell, we took the mean activity across time, and computed a quantile of mean activity relative to all other cells in the session. To classify whether a cell is a place cell, we circularly shuffled the deconvolved signal for each cell 500 times. The same calculation was repeated to compute the spatial information and stability for these shuffled activities and to build a null distribution of spatial information and stability for each cell. We classified a cell as a place cell if its spatial stability and spatial information were greater than 95^th^ percentile of the corresponding null distribution. In other words, a place cell was defined as a cell with both significant spatial information and significant spatial stability (*p* < 0.05 compared to chance). To compute population vector (PV) correlations between a pair of sessions, we built the PV for each spatial bin from the spatial firing activity across all place cells. For cells in the GCaMP channel, we handled inactive cells in two different ways when building the PV: either the spatial firing activity of inactive cells were set to zero, or the inactive cells were excluded from the PV (leaving only the place cells that were active in both sessions). Then, we computed the Pearson correlation of the PVs corresponding to the same spatial bin, and we used the mean correlation across all spatial bins as the PV correlation between a pair of sessions.

## Supplemental information

**Table S1:** Bill of materials for dual-channel Miniscope. The name, vendor, part number and description are shown for each part.

| Name | Vendor | Part number | Description |
| --- | --- | --- | --- |
| Excitation filter | Chroma | ZET488/561x | 488nm and 561 nm dual-band pass |
| Excitation dichroic mirror | Chroma | ZT488/561rpc | Reflects wavelengths between 482-493nm and 555-566nm |
| Emission dichroic mirror | Chroma | T556lpxr | 556nm high-pass |
| Side emission filter | Chroma | ET525/50m | 525nm band pass |
| Top emission filter | Chroma | ET600/50m | 600nm band pass |
| LED | Lumileds | LXZ1-PB01 | high-power LED with 460-480nm dominant wavelength |
| Objective achromatic lens | Edmund Optics | 45-089 | 3mm diameter 6mm FL achromat |
| Achromatic lens | Edmund Optics | 63-691 | 4mm diameter 10mm FL achromat |
| Electro-wetting lens | Edmund Optics | 34-282 | 2.5mm CA A-25H0 Corning Varioptic Variable Focus Liquid Lens |

**Figure S1:**
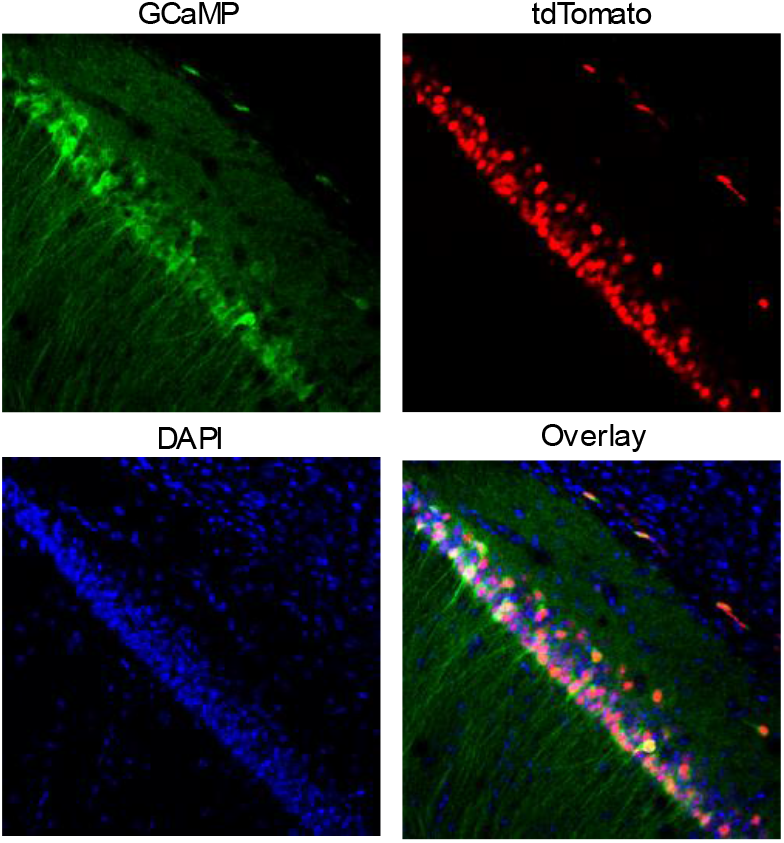
Confocal validation of virus expression. Expression profile of AAV1-hSyn-GCaMP6f-P2A-nls-dTomato virus in hippocampus CA1 is shown.

**Figure S2:**
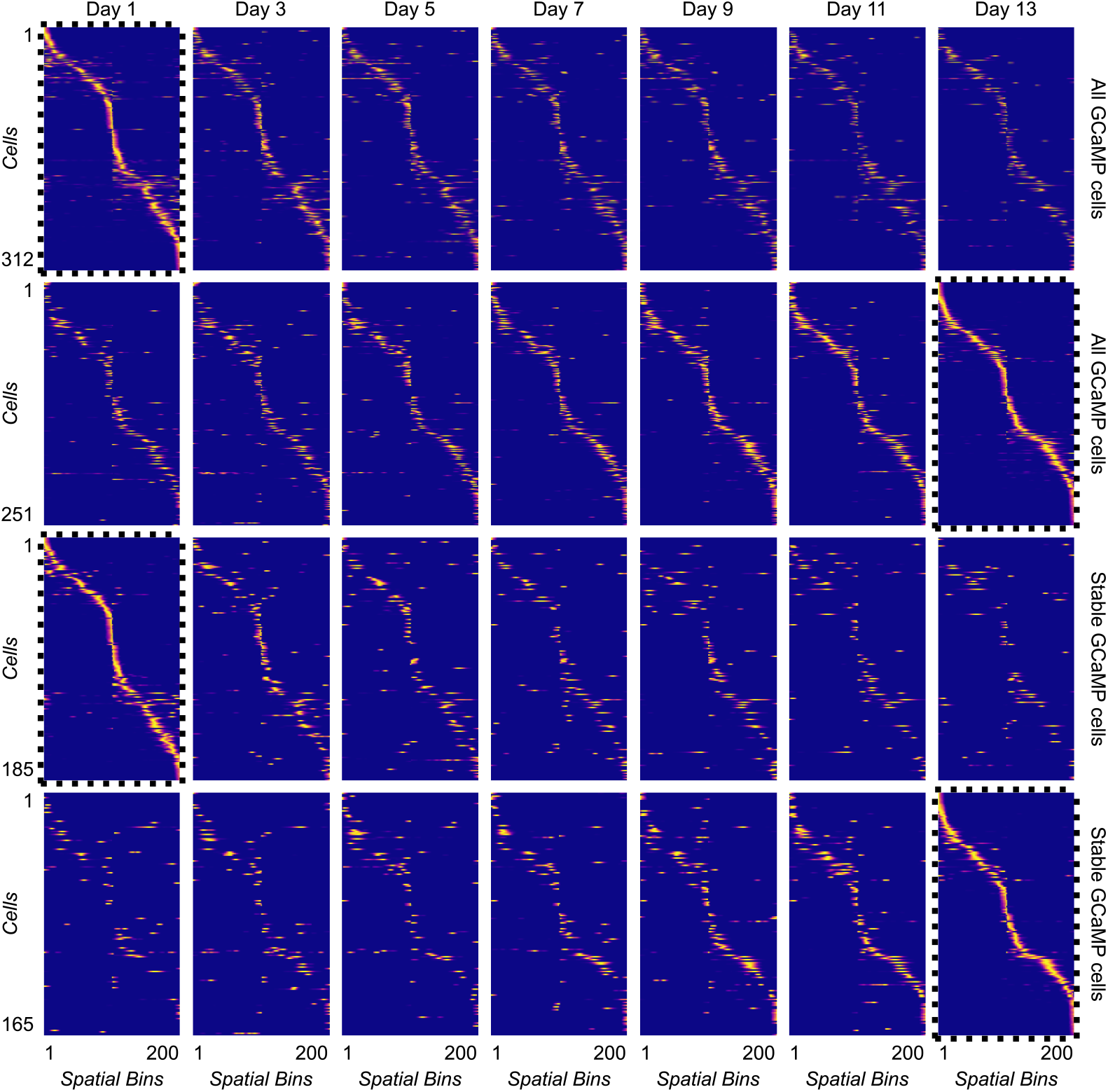
Spatial firing patterns of cells tracked across the full experiment. Each row represents a cell tracked throughout the experiment and each column represents a spatial bin. The average firing rate is shown across 200 spatial bins (100 spatial bins for each running direction). First row: all cells in GCaMP channel are included and sorted according to peak firing position in the first day. Second row: same population of cells as first row but cells are sorted according to peak firing position in the last day. Third row: only GCaMP cells whose corresponding tdTomato landmark can be tracked stably across all 7 recording sessions are included. The cells are sorted according to peak firing position in the first day. Fourth row: same population of cells as the third row but cells are sorted according to peak firing position in the last session.

## Acknowledgements

The authors acknowledge support from following funding sources: TS is supported by CURE Epilepsy Taking Flight Award, American Epilepsy Society Junior Investigator Award, R03 NS111493, R21 DA049568, and R01 NS116357. DA is supported by U01 NS094286-01, and 1700408 Neurotech Hub. DJC is supported by DP2 MH122399, R01 MH120162, Botanical Center Pilot Award from P50 AT008661-01 from the NCCIH and the ODS (Pasinetti PI), One Mind Otsuka Rising Star Award, McKnight Memory and Cognitive Disorders Award, Klingenstein-Simons Fellowship Award in Neuroscience, Mount Sinai Distinguished Scholar Award, Brain Research Foundation Award, and NARSAD Young Investigator Award.

